# Systematic dissection of biases in whole-exome and whole-genome sequencing reveals major determinants of coding sequence coverage

**DOI:** 10.1101/387639

**Authors:** Yury A. Barbitoff, Dmitrii E. Polev, Andrey S. Glotov, Elena A. Serebryakova, Irina V. Shcherbakova, Artem M. Kiselev, Anna A. Kostareva, Oleg S. Glotov, Alexander V. Predeus

## Abstract

Next generation DNA sequencing technologies are rapidly transforming the world of human genomics. Advantages and diagnostic effectiveness of the two most widely used resequencing approaches, whole exome (WES) and whole genome (WGS) sequencing, are still frequently debated. In our study we developed a set of statistical tools to systematically assess coverage of CDS regions provided by several modern WES platforms, as well as PCR-free WGS. Using several novel metrics to characterize exon coverage in WES and WGS, we showed that some of the WES platforms achieve substantially less biased CDS coverage than others, with lower within- and between-interval variation and virtually absent GC-content bias. We discovered that, contrary to a common view, most of the coverage bias in WES stems from mappability limitations of short reads, as well as exome probe design. We identified the ~ 500 kb region of human exome that could not be effectively characterized using short read technology. We also showed that the overall power for SNP and indel discovery in CDS region is virtually indistinguishable for WGS and best WES platforms. Our results indicate that deep WES (100x) using least biased technologies provides similar effective coverage (97% of 10x q10+ bases) and CDS variant discovery to the standard 30x WGS, suggesting that WES remains an efficient alternative to WGS in many applications. Our work could serve as a guide for selection of an up-to-date resequencing approach in human genomic studies.

## Introduction

Next-generation sequencing (NGS) is rapidly becoming an invaluable tool in human genetics research and clinical diagnostics (reviewed in van Dijk et al., 2014). Practical use of NGS methods has dramatically increased with the development of targeted sequencing approaches, like whole-exome sequencing (WES) or targeted sequencing of gene panels. WES emerged as an efficient alternative to whole-genome sequencing (WGS) due to both lower sequencing cost and simplification of variant analysis and data storage (Wang et al., 2013). More than 80% of all variants reported in ClinVar, and more than 89% of variants reported to be pathogenic, come from the protein-coding part of the genome; this number increases to 99% when immediate CDS vicinity is included. Even allowing for the sampling bias, there is an overall agreement that most heritable diseases appear to be caused by alterations in the protein-coding regions of the genome. Given this, WES has dominated the projects characterizing human genome variation as well as clinical applications. The pioneering 1000 Genomes project (Auton et al., 2015) could not statistically characterize many of the rare variants critical to diagnostics of Mendelian disease due to a limited sample size. In an attempt to get a representative picture of protein-coding variation in human population, 6500 WES samples were sequenced during ESP6500 project (Fu et al., 2012). When a much larger reference set of 60,706 WES experiments was compiled and uniformly processed by the Exome Aggregation Consortium (Lek et al., 2016), it dramatically increased the accuracy of allelic frequencies (AF) estimation in general population. This led to a surprising conclusion that up to 90% of variants reported as causative for Mendelian disease in ClinVar database are observed too often in healthy controls to directly cause disease (Lek et al., 2016). The number of available WES experiments is rapidly increasing, and the latest Genome Aggregation Database (gnomAD) collection includes 123,136 WES experiments alongside with 15,496 WGS. Such impressive number of profiled individuals allows a much more thorough look at human coding genome variation, leading to many useful applications such as estimation of selective pressure across protein-coding regions (Cassa et al., 2017).

Originally the enrichment of exonic sequence was done using hybridization on solid high-density microarray, including the proof-of-concept publication (Hodge et al., 2007), as well as first reports of exome use for diagnostics of Mendelian disease (Ng et al., 2009; Ng et al, 2010). The technology has later shifted towards a more efficient solution-based enrichment strategy involving biotin-tagged probes and streptavidin-coated beads (Gnirke et al., 2009; Bainbridge et al., 2010). Alongside with hybridization-based methods, amplicon-based technologies using ultra-multiplexed PCR have also been developed. Among the currently commercially available technologies few belong to the amplicon type (e.g. Agilent HaloPlex and Thermo Fisher Ion AmpliSeq), while others use hybridization capture systems (e.g., Agilent SureSelect, Illumina TruSeq Exome, and Roche SeqCap EZ). Modern applications heavily favor the latter; for example, ExAC and gnomAD only contain data from hybridization-based technologies. The reason for such preference is much greater variability of generated unique fragments in hybridization-based methods, which increases the library complexity, improving the statistical power of variant calling (Samorodnitsky et al., 2015).

Several published studies have concentrated on comparing the performance of different exome capture technologies. The pioneering studies (Teer et al., 2010; Asan et al., 2011) have compared approaches that either have never become commercialized, or have been completely replaced, and often focused on a region substantially smaller than whole exome. With the emergence of commercial exome kits, three major manufacturers - Agilent, Illumina, and Nimblegen (Roche) - have become popular among users, representing the majority of all published WES studies. Early comparative studies have focused on comparison of target intervals of various exome kits, and identified several important biases inherent to WES technology, such as coverage biases in regions with very high or low GC content (Clark et al., 2011; Parla et al., 2011; Sulonen et al., 2011). A later study comprised most hybridization-based capture technologies available at the time (Chilamakuri et al., 2014), and showed specific features of each of the four exome kits, including GC-content bias and differences in the distribution of coverage. However, this and other earlier studies included very limited number of samples, often with large variation of sequencing depth, which may have interfered with consistent platform comparison.

It is often assumed that WGS offers more uniform coverage of CDS regions due to the nature of hybridization-based enrichment process used in WES. Such differences in coverage evenness increase the costs of effective per-base coverage in WES, questioning the overall benefit from using WES instead of WGS. Hence, the issue of WES/WGS comparison has been addressed by several studies that sought out optimal sequencing method to achieve maximum coverage of the protein-coding regions of the genome. One of these included Agilent and Nimblegen (Roche) WES capture technologies, that were compared with the conventional WGS approach in terms of resulting coverage per sequencing read and the efficiency of clinically significant SNV detection (Lelieveld et al., 2015). Similarly to earlier studies (Clark et al., 2011; Parla et al., 2011), it was found that WES achieves similar percentage of well-covered CDS bases only when the average coverage is 2-3 times higher, and with a substantial sequence bias. In several more recent studies, it was repeatedly stated that WGS provides more even and unbiased coverage of coding regions and generates more accurate variant calls (Belkadi et al., 2015; Meienberg et al., 2016; Carss et al., 2017).

Hybridization probes in exome kits are designed based on most comprehensive annotations of human coding sequence available. The part of human transcriptome annotated as coding has been converging, according to GENCODE/Ensembl/RefSeq version history. Manufacturers are also constantly improving their WES solutions, and all commercial providers are currently offering a number of kits with extended UTR coverage and with bait designs updated according to more recent annotations of human genome. Importantly, variant-calling practices are also entering a more mature phase, with Genome Analysis Toolkit (GATK) Best Practices (DePristo et al., 2011; van der Auwera et al., 2013) providing the a comprehensive and well-validated set of recommendations that make variant discovery accurate and reproducible. Hence, in this study we took an effort to make an up-to-date and thorough assessment of the performance of the most recent versions of four widely used exome capture systems (Illumina Nextera Rapid Capture v1.2, Illumina TruSeq Exome v1.2, Agilent SureSelect AllExon v6, and Nimblegen (Roche) SeqCap EZ MedExome) on a cohort of several hundred patients, and compare the performance of WES technologies with the most recent PCR-free WGS, which has been shown to outperform some WES technologies in coverage efficiency (Meienberg et al., 2016). We show that best modern WES platforms allow for efficient coverage and variant discovery in CDS regions and, despite frequent statements, are not dramatically outperformed by WGS. Our study may serve as a guide for selection of the resequencing approach in research and clinical practice of human genetics, and uncovers important determinants of coding sequence coverage in human genome.

## Online methods

### Samples

Peripheral venous blood samples were collected in EDTA from 167 patients with endocrine diseases, hereditary connective tissue disorders, orphan diseases and individuals from the control group. All patients gave informed consent for blood sampling, research, processing of personal data and storage of biological materials. DNA was extracted with QIAsymphony automated station for the isolation of nucleic acids and proteins. All WES samples satisfied the ExAC criterion of 80+% of CDS bases with 20x coverage.

### Exome library preparation

After DNA extraction, we prepared whole exome libraries with Illumina Nextera Rapid Capture Exome, Nimblegen (Roche) SeqCap EZ MedExome, Illumina Truseq Exome, and Agilent Sureselect XT2 V6 technologies.

#### SeqCap EZ MedExome Kit (Roche, USA)

1 mkg of human DNA in 1x Low TE buffer (pH=8.0) was used as a starting material and sheared on Diagenode BioRuptor UCD-200 DNA Fragmentation System to the average DNA fragment size of 170-180 bp. The shearing conditions were as follows: L-mode, 50 minutes of sonication cycles consisting of 30 s sonication and 30 s pause. Library preparation and exome capture were performed using SeqCap EZ MedExome Kit (Roche, USA) following the SeqCap EZ Library SR User’s Guide, v5.1 without modification. DNA libraries were amplified using 7 PCR cycles, and 14 PCR cycles were performed for amplification of enriched libraries. Library quality was evaluated using QIAxcel DNA High Resolution Kit on QIAxcel Advanced System.

#### Nextera® Rapid Capture Exome Kit (Illumina Inc., USA)

Library preparation and exome capture were performed following the Nextera Rapid Capture Enrichment guide v. 15037436 (Illumina Inc., USA) without modifications. 50 ng DNA was used as a starting material and 10 cycles of PCR were performed for pre-enrichment and post-enrichment PCR steps. Library quality was evaluated using QIAxcel DNA High Resolution Kit on QIAxcel Advanced System.

#### TruSeq Exome Library Prep Kit (Illumina Inc., USA)

300 ng of human DNA in 100 μl of 1x TE buffer (pH=8.0) was used as a starting material and sheared on Diagenode BioRuptor UCD-200 DNA Fragmentation System to the average DNA fragment size of 200 bp. The shearing conditions were as follows: L-mode, 45 minutes of sonication cycles consisting of 30 seconds sonication and 30 seconds pause. Shearing results were evaluated using QIAxcel DNA High Resolution Kit on QIAxcel Advanced System. Several (1-10) additional sonication cycles were performed to reach the desired 200 bp DNA fragment size peak, when needed. 100 ng of sheared DNA was used as a starting material for library preparation. Library preparation and exome capture were performed using TruSeq Exome Library Prep Kit following the standard TruSeq Exome Library Prep Reference Guide (Illumina Document # 15059911 v01). Library quality was evaluated using QIAxcel DNA High Resolution Kit on QIAxcel Advanced System.

#### Agilent SureSelect XT2 Library Prep Kit ILM v.6

2 mkg of human DNA in 100 μl of 1x Low TE buffer (pH=8.0) was used as a starting material and sheared on Diagenode BioRuptor UCD-200 DNA Fragmentation System to the average DNA fragment size of 150-200 bp. The shearing conditions were as follows: L-mode, 60 minutes of sonication cycles consisting of 30 seconds sonication and 30 seconds pause. Shearing results were evaluated using QIAxcel DNA High Resolution Kit on QIAxcel Advanced System. Library preparation and exome capture were performed following the SureSelectXT Target Enrichment System for Illumina Multiplexed Sequencing Protocol (Version B5, June 2016) for 3 mkg of starting DNA. Library quality was evaluated using QIAxcel DNA High Resolution Kit on QIAxcel Advanced System.

### Whole-exome sequencing

We used Illumina HiSeq 2500 and Illumina HiSeq 4000 platforms for sequencing. Each exome library was sequenced using 101 bp (HiSeq 2500) or 150 bp (HiSeq 4000) paired-end reads.

### Whole-genome sequencing

For comparison of exome capture technologies with conventional WGS approach, we used several recent samples sequenced at Biobank genome facility (Zhernakova et al., 2018).. WGS libraries were prepared using TruSeq DNA PCR-Free LT Library Prep Kit (Illumina, USA) according to the manufacturer’s protocol. Additionally we used PCR-free WGS data of the Genome in a Bottle (GIAB) consortium (Zook et al., 2017) (Chinese and Ashkenazi trios), as well as several samples publicly available at the NCBI Sequencing Read Archive (SRA) (SRA IDs SRR2098244, SRR2969967, ERR2186302, SRX2798634, SRX2798624). For GIAB samples, we used pre-calculated *Novoalign* BAM files available at the GIAB FTP site (ftp://ftp-trace.ncbi.nlm.nih.gov/giab/ftp/data/). For our own WGS samples and samples downloaded from SRA, we used *bwa mem* v. 0.7.1 for read alignment. All BAM files were narrowed down to the GENCODE v19 CDS regions using *bedtools* (Quinlan and Hall, 2010). We further down-sampled the 300x BAM file for GIAB sample HG001 to obtain 5 separate BAM files with 60x mean coverage or 10 BAM files with 30x mean coverage (on Fig. 3a).

### Whole-exome sequencing data analysis

For all exome and genome samples, bioinformatic analysis of sequencing data was done using a pipeline based on *bwa mem* (Li & Durbin, 2009), *PicardTools* v. 2.2.2 (http://broadinstitute.github.io/picard/) and *Genome Analysis ToolKit* (according to the GATK Best Practices workflow (DePristo et al., 2011; van der Auwera et al., 2013)). Sample genotyping was done in a cohort calling mode using *GATK HaplotypeCaller*. Variant calling was restricted to either bait regions for each technology or the CDS regions (see Results). Variants were filtered using *Variant Quality Score Recalibration* (SNV sensitivity 99.9%, indel sensitivity 90.0%). Annotation and subsequent filtration of variants was done using *SnpEff* and *SnpSift* tools followed by automated correction of reference minor alleles by *RMA Hunter* (Barbitoff et al., 2018). Hybrid selection metrics were calculated using *CollectHsMetrics* tool in the *PicardTools* package. Alignment data visualization was carried out in the *Integrated Genomics Viewer (IGV)* (Robinson et al., 2011).

### Interval file comparison

To analyze the proportion of CDS and UTR sequences covered by each technology’s declared design file, we used the *bedtools* package (Quinlan & Hall, 2010). Reference GENCODE v19 genome annotation (http://gencodegenes.org/) (Harrow et al., 2012) was used for these estimations. Only chromosome located CDS regions of protein-coding genes were used in the analysis. We also used ClinVar database of variants implicated in human disease (build 2018-04-01) to assess coverage of important variant sites.

### Coverage calculation

Modern best practices advise using *GATK* toolset for variant calling, which ignores reads with mapping quality (MQ) less than 10 and reads mapped as duplicate by *Picard MarkDuplicates* utility. Thus, all coverage calculations were done on BAM files with duplicate reads and reads with MQ < 10 removed. Exact coverage calculation pipeline is available at https://github.com/bioinf/weswgs.

### Calculation of coverage evenness statistics

To analyze the distribution of coverage across target regions, as well as between-interval evenness (BIE) and within-interval evenness (WIE, or coverage smoothness), we used a combination of *bedtools* package and custom scripts in *bash* and *Python* (available at https://github.com/bioinf/weswgs). To collect normalized coverage profiles for each platform, BAM files were converted to a bedGraph format using *bedtools* (Quinlan & Hall, 2010). Next, the bedGraph file was intersected with the CDS regions according to GENCODE v19 genome annotation or the declared target regions for each technology. To calculate the coverage evenness and profiles of per-base normalized coverage we used the resulting bedGraph files to obtain fractions of bases having normalized coverage of at least N with N ranging from 0 to 3 with step 0.01. Evenness score was calculated as described (Mokry et al., 2010).

To calculate the between-interval evenness (BIE), mean sequencing depth was calculated for each interval. These coverage values were then processed similarly to per-base coverages. Between-interval evenness (BIE) measure was calculated similarly to the OE from the profiles of normalized mean coverages of individual intervals.

For calculation of the within-exon coverage distribution, all intervals having an average coverage of more than 10x in a sample were then divided into 100 bins of equal length. We then calculated normalized (divided by the mean coverage of a fragment) coverage in each bin. Then, mean coverage at each bin across all of the intervals was calculated for each sample. Within-interval evenness (WIE, or smoothness) was defined as the area under within-interval normalized coverage curve (restricted to the maximum value of 1):

where *i* is the bin number (relative distance within the interval with step of 0.01), and *x*_*i*_ is the normalized coverage in this bin.

### Variant calling performance analysis

To calculate the allele ratio distribution and the distribution of total and low genotype quality (lowGQ) variants, we used the VCF file resulting from the cohort genotyping of samples, and scripts written in Python (available at https://github.com/bioinf/weswgs). To calculate the mean coverage of variant sites depending on the GC-content of variant site neighborhood we selected 180452 known variants from the ClinVar database of clinically significant variants, and divided these variant sites into 10 equal groups depending on the GC-content (calculated by *bedtools nuc*) of the region 50 bp up- and downstream of the variant. We then calculated the read depth at all resulting variant sites using *bedtools multicov*.

### Modelling coverage biases

To construct a model of per-interval normalized coverage, we have calculated mean normalized coverage of each individual CDS region in all samples sequenced with a particular technology, as well as the standard deviation of the mean. We also calculated multi-mapping fraction (MF) by subtracting mean coverage after filtering by mapping quality (MQ > 10) from mean coverage before such filtering. We next predicted the amount of base-pairs with low (< 10x) coverage by sampling the value of mean normalized interval coverage and calculating per-base normalized coverages by multiplication of sampled mean interval coverage value and a WIE profile for a given interval. This procedure was repeated for a range of exome average depths (20-200x). To obtain a list of intervals that systematically or randomly have poor sequencing coverage for each technology, we statistically evaluate the difference between the mean normalized coverage of each CDS region and the threshold value using one-sample *t*-test with Holm-Bonferroni FDR correction. To assess the reproducibility of coverage biases, we calculated mean pairwise correlation of vectors of per-interval normalized coverages across all samples sequenced with a particular platform (or all platforms in the study). To assess the relative importance of different variables for prediction of normalized per-interval coverage, we fitted a simple linear model describing mean per-interval coverage depending on GC-content, interval length, multi-mapping fraction, and a binary variable indicating inclusion of the interval into the exome design. Predictor importance was analyzed by the value of lmg statistic using *relaimpo R* package (Gromping, 2006). To use random forest classification model for prediction of poorly covered regions, we constructed a binary variable indicating a normalized coverage < 0.1 and used it as a target for prediction by *randomForest R* function with the predictors described above.

### Estimation of the ExAC variant site density

Exome Aggregation Consortium (ExAC) variant calls (v. 0.3.1) (Lek et al., 2016) were used to make statistical assessment of variant density. Each exome interval was annotated with the number of ExAC variant sites that fall inside this interval using *bedtools intersect*. Next, variant counts were transformed to per-nucleotide variant site density, and the resulting dataset was used for sampling procedures.

### Data availability and scripts

All statistical analyses were carried out using *R*. All graphs were plotted using *ggplot2* (Wickham, 2016) and *cowplot* packages. Scripts for data analysis and figures, as well as the processed data, can be found at https://github.com/bioinf/weswgs.

## Results

### WES bait design efficiency

Over the recent years, advances in human genome analysis have led to a dramatic convergence of genome annotations by major consortia (GENCODE, RefSeq, Ensembl). Our analysis shows that latest GRCh37-based modern genome annotations (RefSeq, Ensembl v75, GENCODE v19) agree on ~ 33.324 Mb out of 35.453 Mb potential chromosomal CDS sequences (94.0%). Such annotation convergence could cast significant improvements onto the quality of bait design for exome sequencing. Hence, we first compared the designs of all modern whole-exome technologies used in the study to assess whether exome designs have similarly converged. We evaluated the intersection of target regions and CDS intervals (based on GENCODE v19) covered by each design and proportion of in-CDS clinically significant variants (as denoted in the ClinVar database) (Table 1). The results show that all platforms cover most of CDS regions (with > 90% CDS bases included in the bait intervals), with Illumina designs being the most comprehensive (99.1% of CDS bases). Declared SureSelect bait intervals are much larger than the ones provided by the other platforms (60.5 Mb vs. ~ 45 Mb), reflecting the compound size of the empirical regions covered in an average experiment. It is noteworthy that SureSelect intervals cover the smallest fraction of CDS bases compared to other technologies (Table 1). We also discovered substantial differences in the amount of ClinVar variants (based on ClinVar build 2018-04-01) covered by each design. The coverage of known pathogenic variants (according to ClinVar) is nearly identical to the percentage of CDS intervals covered, reflecting the fact that all manufacturers ensure thorough coverage of known disease genes. All kits included in the study do not feature extended UTR coverage and include ~20% of GENCODE v19 UTR regions. It is important to distinguish factual coverage from interval design: due to the nature of bait enrichment, even the CDS regions that are not technically included in the designed intervals could be effectively covered.

**Table 1.**
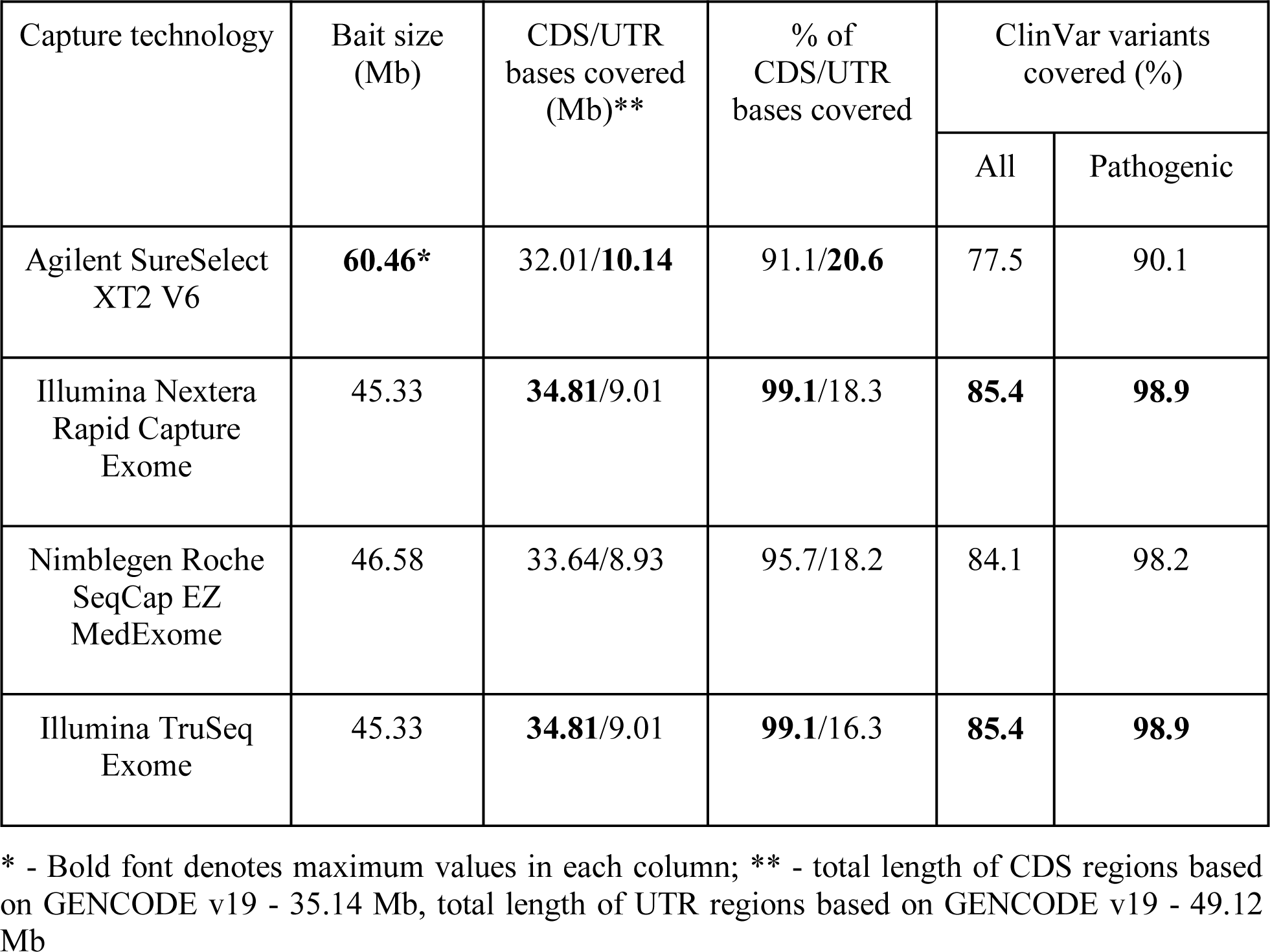
Comparison of CDS coverage by target regions.

### Coverage evenness analysis within and between CDS regions

We then turned to characterize the efficiency of CDS interval coverage by the listed exome capture technologies. It is important to note that most modern variant calling tools ignore reads with mapping quality (MQ) less than 10 and reads marked as PCR or optical duplicates; thus, such reads were removed when calculating coverage. Irrespective of the platform, all samples showed ~50-70% efficiency of target enrichment and a similar distribution of sequencing depths across our WES dataset (Fig. 1a,b), corresponding to 38 ± 5 fold enrichment of target regions (Supplementary Fig. 1). Interestingly, we observed a weak trend showing that libraries having higher depth of sequencing tend to show less efficient exome enrichment. The strength of the trend depends on the particular technology: for SureSelect and TruSeq Exome kits the trend is almost absent (R^2^ = 0.034 and R^2^ = 0.026, respectively), while for MedExome and Nextera Rapid the dependence is much more pronounced (R^2^ = 0.360 and R^2^ = 0.815) (Fig. 1c).

**Figure 1.**
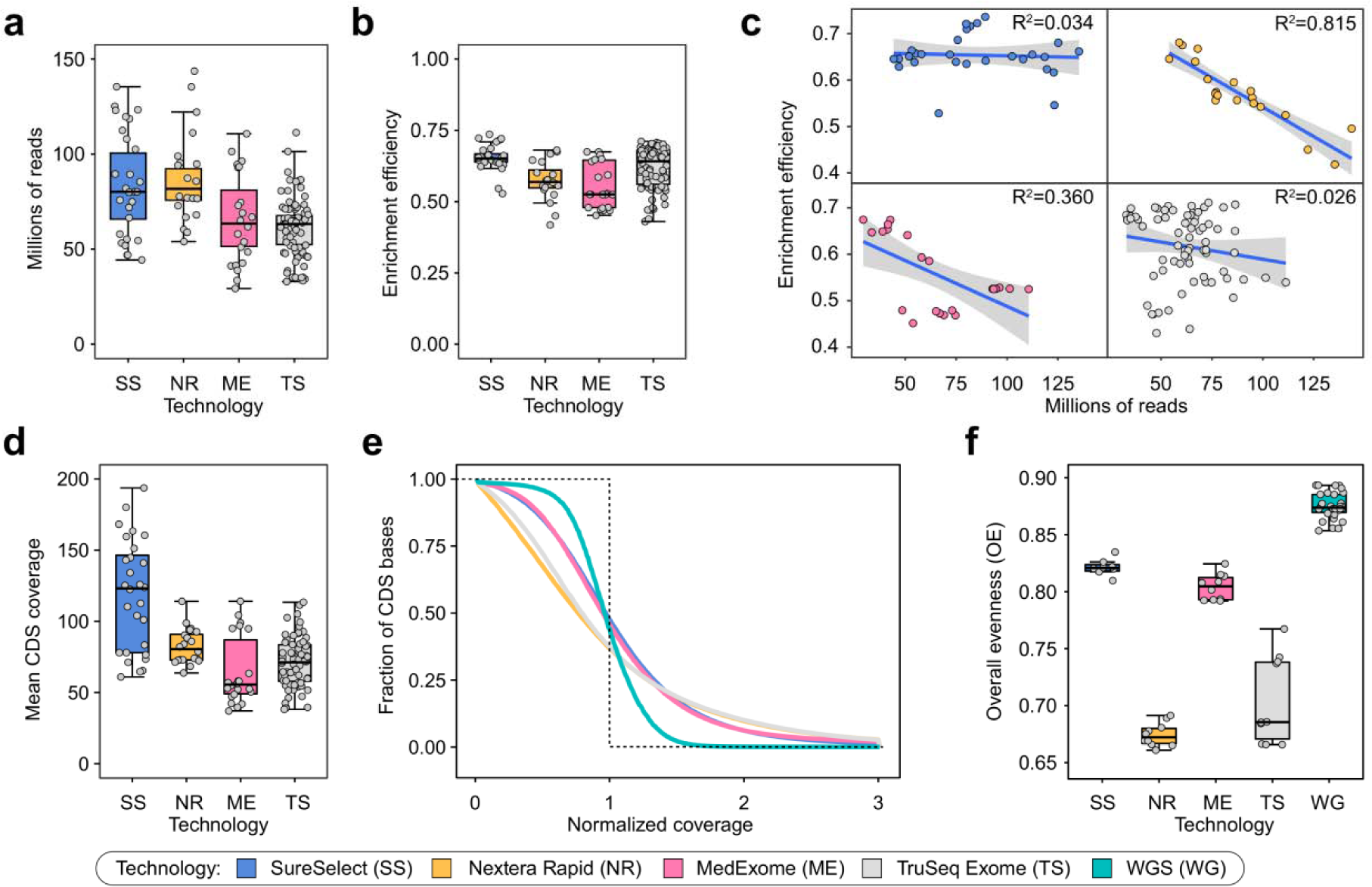
Coverage of target regions across WES and WGS samples. **(a, b, d)**. Total read depth (a), target enrichment efficiency (b) and mean CDS coverage (d) for all samples for each platform **c.** Enrichment efficiency dependence on total read depth. Lines are linear regression fits with confidence intervals indicated as grey envelopes. **e.** The distribution of the normalized coverage for all WES technologies compared to WGS. Dotted line represents ideal case baseline, i.e. all bases covered at mean value. **f.** Overall evenness scores, calculated as described in Mokry et al., 2010, for all four WES technologies and WGS. For plots (e, f) a subset of 10 samples with similar mean coverages was selected for all WES platforms.

Mean coverage of CDS regions in our dataset was comparable among different capture technologies (~ 70x), with exception for SureSelect, that had mean coverage of ~ 120x (Fig. 1d). We then calculated profiles of normalized coverage across CDS bases (Fig. 1e). In order to characterize the overall evenness of CDS coverage (OE), we have used the score developed by Mokry et al (Mokry et al., 2010; see Methods). Normalized coverage profiles and OE scores showed that both Illumina kits perform significantly worse than SureSelect and MedExome, while all exome platforms provided less even coverage than WGS (Fig. 1e-f).

To dissect potential sources of coverage bias we defined two possible components of coverage evenness: coverage distribution between different CDS regions (between-interval evenness, BIE), and uniformity of coverage within individual intervals (within-interval evenness, WIE). The latter type of coverage unevenness is inherent to WES platforms; hence, we first questioned whether it explains the difference between WES and WGS in the OE scores. Indeed, visual inspection of coverage profiles on individual CDS regions suggests that exome platforms highly vary in WIE (Fig. 2a). To more accurately assess the observed differences, we calculated average profiles of relative coverages and WIE scores for all CDS regions (Fig. 2b). We found that WIE scores are well correlated with the OE (Fig. 2c), however, WIE does not completely explain differences observed in Fig. 1f. Similar results were obtained by calculation of WIE profiles across CDS intervals including flanking regions and with exon length stratification (Supplementary Fig. 2). Importantly, WGS did not show any noticeable within-interval unevenness, confirming that such type of coverage bias is specific to WES platforms. As our results suggest WIE is not the only source of increased coverage bias in WES, we next calculated profiles of relative mean interval coverage across all CDS regions to estimate BIE (Fig. 2d). We observed that, while WGS generally performed better than WES, exome platforms showed a distinct pattern of between-group differences (Fig. 2e) that explains the discrepancy between OE and WIE scores, implying that the overall coverage evenness is the product of both BIE and WIE.

**Figure 2.**
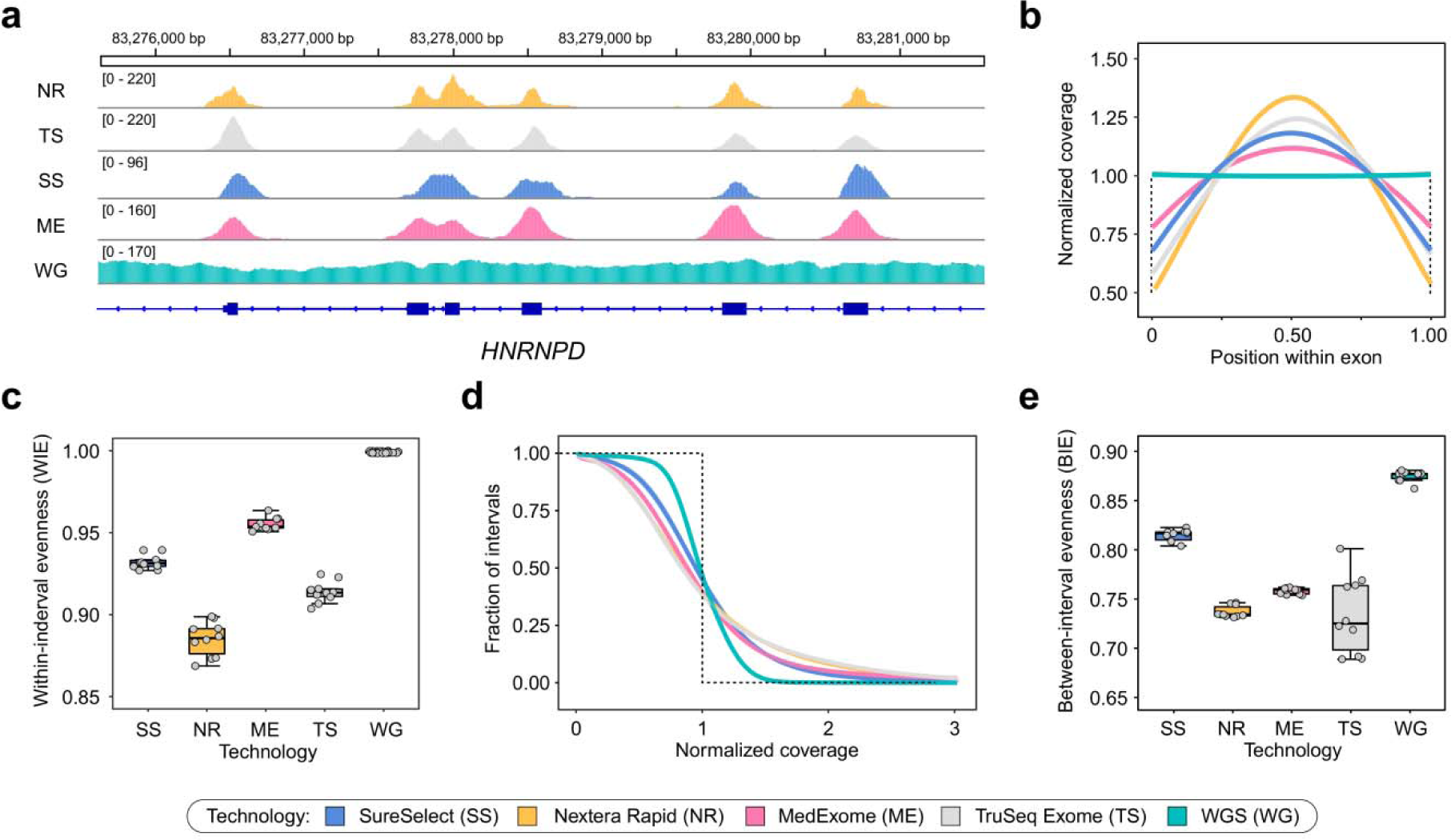
Different technologies exhibit specific patterns of coverage within exons and differ coverage distribution within exons. For all plots, a subset of samples was used as described earlier. **a.** Example of sequencing coverage patterns across exons of the *HNRNPD* gene. Shown are selected samples with similar mean CDS coverage. **(b, c)**. Within-interval coverage evenness (WIE). **b.** Distribution of relative coverage from the start to the end of target interval, averaged over all CDS regions. **c**. WIE scores calculated from distributions in (d) (see Methods for more details; all capture technologies differ in pairwise U-test with Holm-Bonferroni FDR correction (adjusted p-value < 0.001)). **(d, e)** Between-interval coverage evenness (BIE; see Text for definitions). **d.** Distribution of normalized mean coverage across CDS intervals. As in Figure 1, dotted line represents ideal case baseline. **e.** Between-interval evenness scores derived from normalized coverage curves in (a).

### Relative importance of coverage bias determinants in WES and WGS

To more thoroughly characterize the capabilities and limitations of resequencing approaches, we have constructed a model to predict sequencing coverage of CDS regions that accounts for both between- and within-interval evenness (see Methods). Quite surprisingly, model-based prediction of the amount of bases covered at less that 10x at different mean coverages showed that all platforms, including WGS, have a certain amount of bases that are not covered at the required depth even at 200x average coverage (Fig. 3a). For common 30x WGS samples, 788 kbp of CDS sequences are covered less than 10x (with 407 kbp covered < 10x at 200x mean coverage); for SureSelect, the best WES platform, 1180 kbp are predicted to have low coverage at 100x, and 970 kbp - at 200x). These results suggest that there are certain sources of reproducible coverage bias for both WES and WGS. We have set out to explore exactly how reproducible are these coverage biases, and what is the relative importance of different sequence features for the efficient coverage of CDS regions and variant discovery in exome sequencing.

To this end we evaluated the correlation of per-interval mean coverages across all samples for each technology. As seen from Fig. 3b, coverage bias in both WES and WGS has a systematic component. Unexpectedly, WGS has shown a more reproducible coverage bias when compared to WES. Among different WES platforms, SureSelect had the lowest reproducibility of coverage across CDS regions, and the two Illumina technologies had significant cross-correlation, suggesting that our estimates reflect specific features of capture process and bait design (Fig. 3b).

We then turned to dissect specific covariates that affect CDS coverage in WES and WGS. We first analyzed the variation in sequencing depth across regions with different GC-content, as GC-content has been referred as a major source of coverage bias in WES (Clark et al., 2011; Meienberg et al., 2017). We calculated normalized coverage at ~180,000 ClinVar variant sites divided into 10 deciles dependent on GC-content of 100 bp variant vicinity, and found that both Nextera Rapid and TruSeq Exome capture kits performed much worse than the others in GC-rich regions and better in the AT-rich ones (Fig. 3c). Among all four WES technologies, MedExome and SureSelect showed the best results with almost no dependence of read depth at variant site on the GC-content of the surrounding region. We also discovered a slight decrease in mean sequencing depth in GC-rich regions for WGS libraries. Overall, our analysis suggests that GC-bias is not a significant factor for best WES platforms and WGS.

We then investigated another plausible source of coverage bias, namely, mappability limitations in short-read sequencing technologies. CDS regions are often considered unique and non-repetitive; though several examples of large repeated CDS elements have been noted (Larson et al., 2015). Curiously, we noticed that for some genes there is a substantial decrease in read depth after exclusion of reads with low mapping quality (MQ). We conservatively define multi-mapping fraction (MF) as the proportion of sequencing coverage that results from reads with MQ = 0. We then calculated MF for each exome base-pair and for individual CDS regions, and analyzed the amount of bases or intervals with high MF (for interval-level analyses, we focused on intervals with MF > 0.4, as this threshold generated 452 kb of sequences of interest, nearly matching the numbers observed in coverage model analysis (Fig. 3a)). On average, exome kits had more bases with higher MF (and, in particular, MF = 1, i.e. all coverage resulting from reads with zero mapping quality) than WGS (Fig. 3d). Strikingly, we found virtually no dependence of the amount of bases covered by multi-mapping reads on read length (Supplementary Fig. 3); and the difference between different WES platforms and WGS appears to be explained mostly by insert length of the sequenced fragment (Supplementary Fig. 4). Overall, ~500 kbp of CDS sequences have MF = 1 even in WGS samples, suggesting that coverage limits for WGS arise mostly from mappability issues.

We next questioned whether a substantial proportion of CDS regions with low normalized coverage in WES samples are simply not targeted by the capture probes. To address this issue, we overlapped regions with normalized coverage significantly lower than 0.1 (or 0.2) (see Methods) with the bait design files, and calculated what fraction of poorly covered bases do not overlap target regions of exome design. Our analysis showed that for most platforms a large fraction of poorly covered bases falls into non-targeted regions of the exome (Fig. 3e). It is also apparent that best WES platforms (SureSelect and MedExome) are almost identical to WGS in the number of targeted bases that are poorly covered in all samples.

Finally, to compare the relative importance of different factors influencing CDS coverage, we fitted a linear regression model to predict normalized per-interval coverage depending on GC-content, interval length, multi-mapping fraction, and inclusion of the interval into the exome kit design. Analysis of the model showed that for best WES platforms, i.e. SureSelect and Roche MedExome, multi-mapping fraction and inclusion are the most important predictors of normalized coverage; however, GC-content is the major determinant of coverage for both Illumina kits. We have also used random forest classifier to assess relative importance of the same factors to predict binarized relative coverage thresholded at 0.1. While MF had higher relative weight, the results allow for the same conclusions: multi-mapping fraction and exome kit design remain by far the most influential factors (Fig. 3f).

Among regions with high MF, we found ~2000 exons corresponding to more than 500 genes, including known disease genes and cancer driver genes (Supplementary Table 2; Fig. 4a). Enrichment analysis of these genes over canonical pathway list from MSigDB showed significant overlaps with diverse immune system-related gene sets (Fig. 4b). This result is not surprising given that immunity-related genes are among the most duplicated gene families in the vertebrate genomes (Nei et al., 1997). Our analysis only accounts for chromosomal parts of the 1000 genome assembly (also known as “b37”), which is most often used for variant calling. Including alternative contigs (totaling 3-4 Mbp depending on genome version and annotation, Fig. 4c) would certainly increase the ambiguity, especially when not using alt-aware alignment and variant-calling tools. Importantly, current genome annotations also contain up to ~40 kbp of coding sequence in primary extrachromosomal scaffolds, which are not covered by any of the current WES platforms.

**Figure 3.**
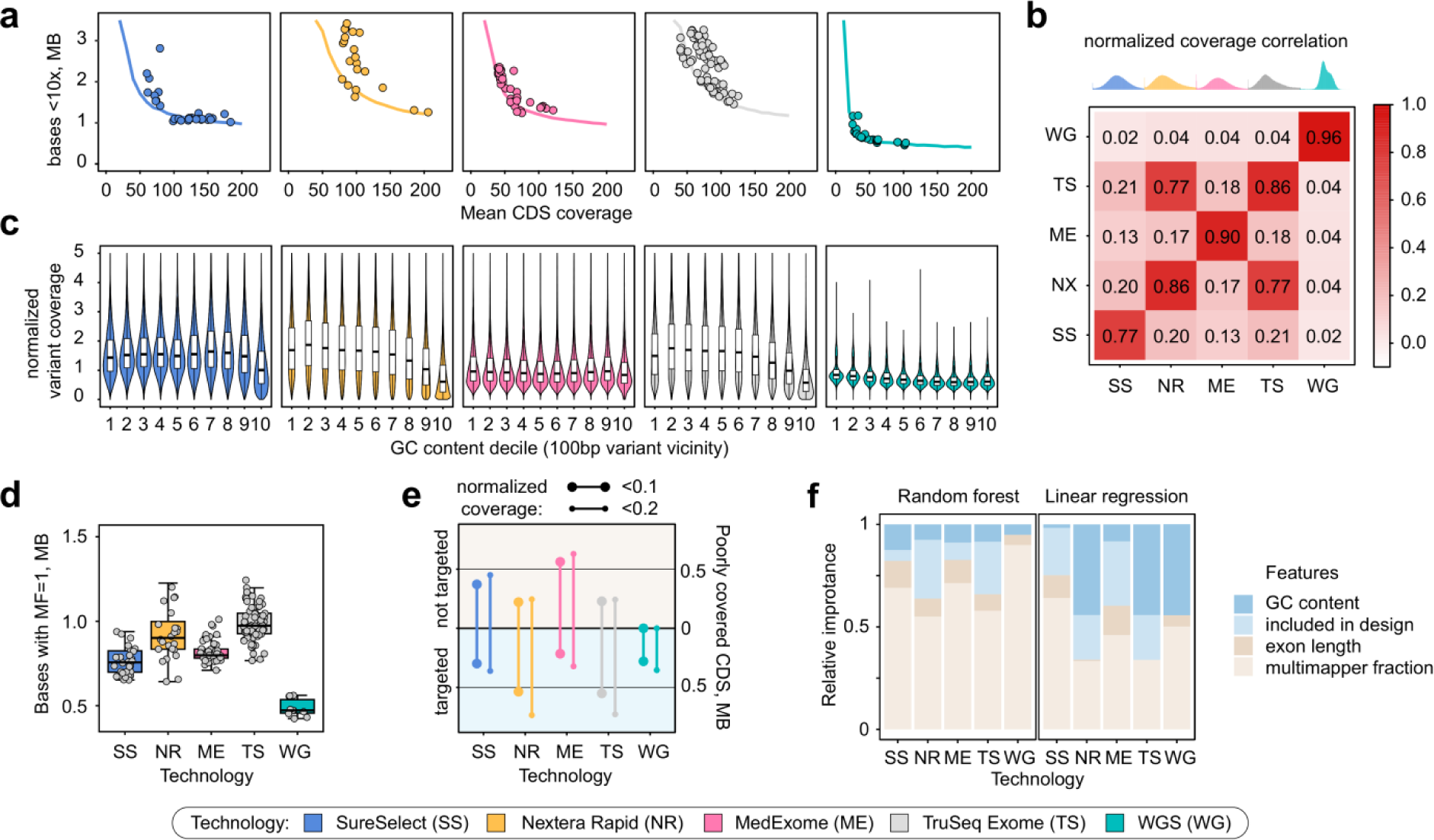
Modeling of CDS coverage identifies key determinants of coverage evenness. **a.** A model based on normalized coverage patterns suggests existence of coverage limits for each technology. Solid lines correspond to model predictions of the amount of bases covered < 10x depending on mean CDS coverage. Dots are samples analyzed in the study. **b.** A heatmap showing average correlation between mean coverages for each exon. Distributions on top are those of per-interval normalized coverages. **c.** GC-bias of coverage at variant sites with different GC-content of 100 bp vicinity (median GC-content in each bin: 0.33, 0.38, 0.42, 0.45, 0.49, 0.53, 0.57, 0.61, 0.65, and 0.71). **d.** Comparison of the amount of CDS bases covered only by multi-mapping reads for each technology. **e.** Total length of targeted and not targeted CDS regions with reproducible low (< 0.1 or < 0.2 average) normalized coverage. **f.** Relative importance of different interval features for prediction of normalized coverage using binary classification (random forest, left) or linear regression (right) (see Methods for details on importanc calculation).

**Figure 4.**
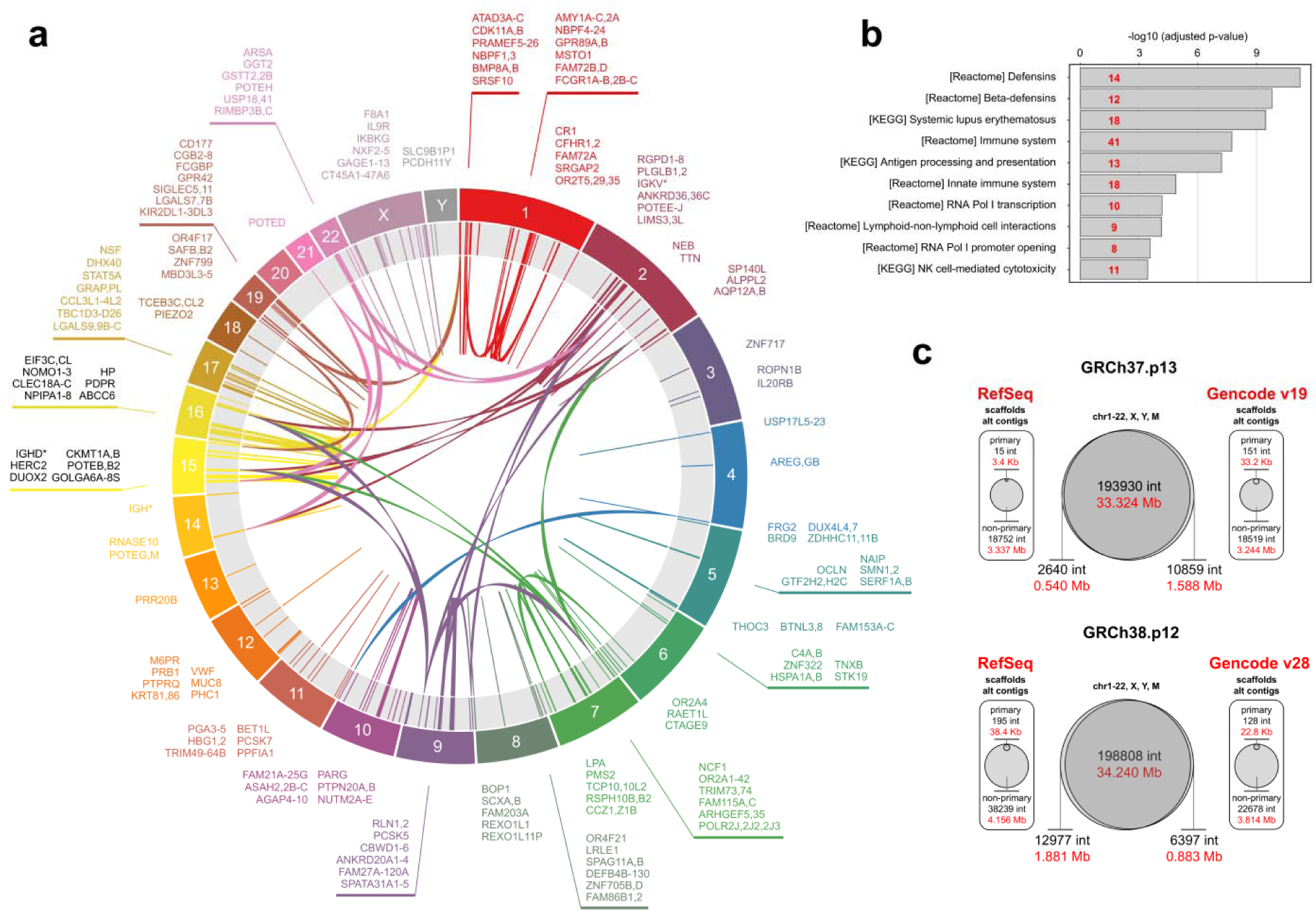
Summary of repetitive human CDS regions inaccessible by current WES and WGS technologies. **a.** Circa diagram showing cross-mappability of CDS regions. Only a subset of clinically relevant genes is shown to decrease diagram complexity. **b.** MsigDB enrichment analysis of genes with CDS regions having MF > 0.4 using canonical pathways (CP) list. Top-10 significant hits are shown. Numbers in red indicate the number of genes in each overlap. **c.** Comparison of modern GRCh37-(top) and GRCh38-based (bottom) genome annotations by RefSeq and GENCODE. Middle, diagrams showing the overall agreement between annotation on chromosomal coding sequences. Summary of extrachromosomal coding sequences for each annotation source is shown aside of the central diagram.

### Variant calling performance on WES and WGS data

In order to see how the observed coverage limitations translate into our ability to detect variation, we have compared the number of variants discovered within CDS for each of the samples in our dataset to summarize the performance of resequencing technologies. We found that for all platforms the numbers of discovered in-CDS variants is approximately the same (Fig. 5a, upper panel), while the number of variants inside CDS that fall within targeted regions is in good correlation with the overall size of the CDS regions covered by each design (Fig. 5a, lower panel). The same proportion is true for the overall amount of variants detected inside the bait regions). The amount of variants with low genotype quality was significantly higher for both Illumina technologies and the highest for the Nextera Rapid kit, while best exome platforms did not differ from WGS in variant call quality (Fig. 5b). Similar results were observed for small insertion-deletion variants (indels); however, WGS have generated slightly fewer lowGQ varians than any of the WES platforms (Fig. 5c-d). Overall, it is very important to note that restriction of variant calling to the bait regions decreases the power of variant discovery in WES, which is otherwise comparable to such of WGS.

**Figure 5.**
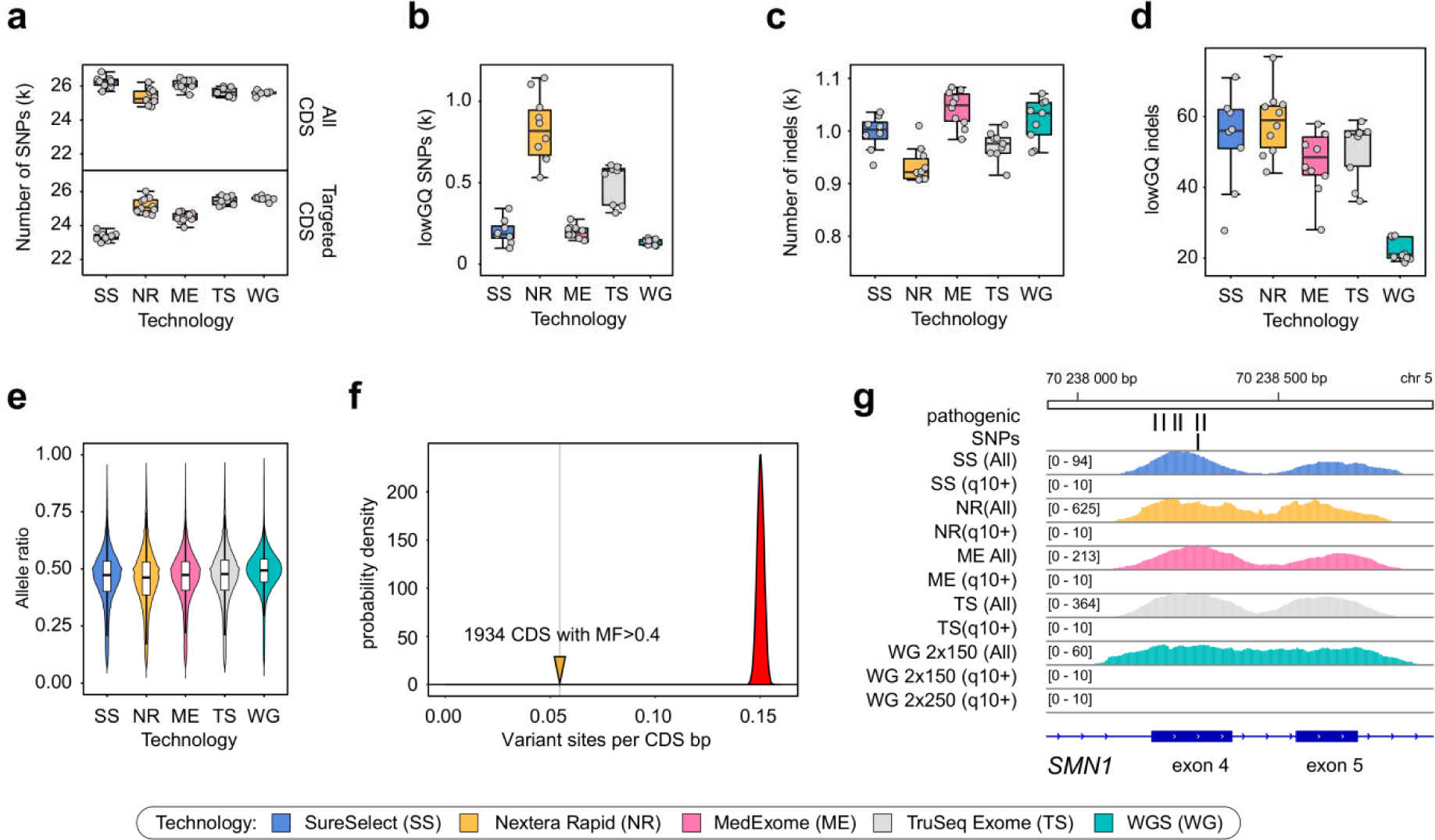
Variant calling biases of 4 exome capture technologies and WGS. **a.** Total number of variants detected inside GENCODE v19 chromosomal coding sequences (all CDS, upper panel) and within targeted CDS regions (targeted CDS, lower panel). **b.** Number of variants with low genotype quality (lowGQ) according to the *GATK GenotypeRefinement* annotation. **(c-d).** Number of all called **(c)** and low-genotype quality **(d)** indels. **e.** Allele ratios at heterozygou variant sites. **f.** Per-nucleotide density of ExAC variant sites in regions with high fraction of multi-mapping coverage (solid line) compared to the distribution of expected variant site density calculated from random subsets of CDS regions (see Methods for details). **g.** Example of the unmappable CDS region in the exons 4-5 of the *SMN1* gene containing several well-established pathogenic variants. Two coverage tracks (including reads with low MQ and excluding these reads) are shown for each technology.

Allele bias is often considered one of the major determinants of poor variant quality in WES samples. To address this issue, we then assessed the allele ratios at heterozygous variant sites. We found no difference between allele ratio distributions in these samples (Fig. 5e) though for Nextera Rapid the distribution is more heavily-tailed with more variant sites having greater coverage of the reference allele. We also calculated the allele bias (AB) ratio characterizing the median amount of reads supporting reference allele in heterozygous variant sites. The AB estimate was found to be ~ 0.53 for MedExome, SureSelect, and TruSeq, while for Nextera Rapid the number was somewhat greater (~ 0.55).

We also statistically assessed the effect of mappability limitations on variant discovery. To this end we have calculated average variant site density (based on ExAC dataset) in CDS regions with MF > 0.4, as well as similar density in 100000 sampled sets of 1934 CDS regions with MF < 0.4. We found that regions with large proportion of multi-mapping coverage have much fewer reported variant sites per nucleotide (Fig. 5f). This result confirms that mappability is an important determinant of sequencing coverage that substantially affects variant discovery. A profound example of a nearly inaccessible CDS region with high clinical relevance are the *SMN1* and *SMN2* genes, mutations in which cause spinal muscular atrophy (SMA) - a fatal neurological disorder with an early age of onset. Indeed, we found that for seven out of eight exons (harboring several well-established pathogenic variants, e.g. rs104893934, rs397514518, rs104893933) inside SMA genes all coverage results solely from reads with zero mapping quality in both WES and WGS (Fig. 5g) (including 2×250 bp WGS). Consistently with these observations, no variants are characterized in these regions in gnomAD exomes and genomes (http://gnomad.broadinstitute.org/gene/ENSG00000172062).

## Discussion

Despite the ready availability of NGS methods, modern large-scale sequencing projects studying human rare diseases and population-scale variation are facing a difficult choice. The often heated “genome vs. exome” debate is complicated by difficulty of estimation of indirect costs of each method. Most sources agree that WGS is 2 to 3 times more expensive, but the number changes a lot between different centers. However, perhaps more importantly, there are few criteria useful to compare method efficiency. The efficiency of the method could indirectly evaluated using percentage of successfully diagnosed cases for Mendelian diseases; most such studies report modest improvement in diagnostic rates (Wright et al., 2018). Other studies have aimed to compare the performance of different WES kits with each other and with WGS more directly, using coverage and variant identification statistics. However, due to the constant improvement in exome kit design and standardization of variant calling procedures, these studies quickly become outdated. Furthermore, low number of reported samples have hampered the use of advanced statistical approaches that would allow to carefully address sample-to-sample variation necessary for such comparison. In this study we have leveraged a unique exome and genome dataset in an effort to provide a universal framework for an unbiased evaluation of modern WES and WGS.

We have found that while all WES technologies provide reasonable enrichment efficiencies, modern SureSelect and MedExome platforms offer substantially more even coverage than both solutions by Illumina (Fig. 1). It has been reported previously that WES provides much less even coverage than WGS (Lelieveld et al., 2015; Meienberg et al., 2017). Our results confirm these statements (Fig. 2e-f); however, a more accurate look at different sources of coverage unevenness suggests that, at least in part, this difference results from within-interval unevenness (Fig. 2) that can be mitigated by increasing sequencing depth. Importantly, modelling of coverage distribution shows that all platforms (both WES and WGS) have significant amounts of CDS bases that are effectively uncovered at any sequencing depth (i.e., at least 407 kb for WGS and 960 kb for best WES; Fig. 3a). This result contradicts earlier statements (Meienberg et al., 2017); however, the reason for such discrepancy is explained by exclusion of reads with zero mapping quality in our analysis pipeline. Variant calling software does not consider reads with low mapping quality; hence, such reads should be omitted in coverage analysis.

Mappability limitations of short-read sequencing technologies render 478 ± 37 kb (for WGS) and 751 ± 34 kb (for best WES) of CDS regions unreachable for sequencing technologies. The problem of low-mappability regions is known; for some of the genes with poor mappability, complex statistical methods have been proposed to determine genotype likelihoods (Larson et al., 2015). However, the mappability issue is often overlooked or considered insignificant for coding regions despite the fact that numerous clinically relevant regions are effectively unmappable (Fig. 4a, Supplementary Table 2), including well-characterized Mendelian disease genes (e.g., *SMN1/SMN2*, Fig. 5g). Statistical analysis of relative predictor importance suggests that, contrary to popular belief, mappability and bait design are the most important determinants of low coverage in WES samples. On the other hand, GC-content, which is usually considered as a major source of coverage bias (Clark et al., 2011; Meienberg et al. 2017; Carss et al., 2017), virtually does not affect coverage for well-designed WES kits or PCR-free WGS (Fig. 3f).

It is important to note that variant calling for WES samples should not be restricted to targeted intervals and should rather include targeted intervals, CDS regions, UTR sequences and bases flanking CDS to improve the power of variant discovery (Fig. 5a-b). Overall, our modeling suggests that a WES sample sequenced with a common 100x average depth will provide significantly poorer coverage of only ~400 kb of CDS compared to a common 30x WGS sample, i.e.in ~1% of coding regions. These predictions are in good concordance with our analysis of variant calling results inside CDS regions (Fig. 5). Best WES platforms are virtually indistinguishable from WGS in both overall number of in-CDS variants discovered and fraction of low genotype quality variants, with WGS showing slightly better performance only for indels. Despite the fact these numbers do not directly estimate each technology’s sensitivity and specificity, they reflect absence of noticeable systematic differences between WES and WGS. A big limitation of all short-read sequencing technologies is their inability to accurately characterize complex structural variants, the problem which will only be solved with newer sequencing approaches based on long reads.

A recent review by Wright et al. (Wright et al., 2018) suggested that WGS is more efficient than WES only by 2% of diagnosis rates on aggregate. Our observations that WGS allows for more efficient coverage of only 1% of exome compared to best WES platforms, as well as the fact that only a small fraction of reported ClinVar pathogenic variants are not targeted by exome kits, are in concordance with these estimates (and, at the very least, imply lack of dramatic increase in diagnosis rate for WGS over WES). Indeed, WGS allows for more accurate mapping of large indels (Carss et al., 2017) and repeat expansions (Bahlo et al., 2018); however, moderate rates of NGS-based diagnostics likely result from several other notable factors, such as lack of biological understanding of variant pathogenicity (Sawyer et al., 2016), inability to characterize redundant genome regions with NGS methods (Fig. 3, see Results), and inefficiency of modern techniques in detection of several important variant classes (Biesecker and Green, 2014), including complex structural variants (Sedlazeck et al., 2018). The latter two reasons demand new technical approaches, while the first requires deeper biological understanding of genome function. In many cases, annual re-analysis of undiagnosed samples using new biological data improves diagnosis rate (Nambot et al., 2018).

Overall, several lines of evidence indicate that WES remains an excellent alternative to WGS in research and clinical applications. Moreover, current WES technologies can be further improved in several ways: first, support for longer insert sizes would decrease the impact of both mappability and WIE; second, inclusion of all chromosomal and especially non-chromosomal CDS regions would make coverage much more comprehensive; and third, better probe design and improvement of hybridization process would alleviate remaining unevenness resulting from GC-content or other sequence-based determinants. In fact, the most recent WES solutions (e.g., produced by Illumina in conjunction with IDT) are reported to perform substantially better than NR or TS kits analyzed in this work, making WES samples approach WGS in terms of coverage distribution and eventually minimizing the diagnostic gap between WES and WGS approaches. Obviously, the inherent limitations of short-read sequencing technologies decrease the power of modern clinical diagnostics much more dramatically than the differences between WES and WGS. Hence, in a more broad perspective, new technical approaches (based on long read single-molecule sequencing) are needed to provide substantially more power for discovery of genetic variants in humans, as well as for diagnosis of rare disease.

## Acknowledgements

We thank Anna Shuvalova and Olga Romanova for help in library preparation. This research was done using equipment of Biobank of the Research Park of SPBU. The research was supported by Russian Science Foundation (grant no. 14-50-00069) and CAF Charity Foundation. We also thank Resource Center “Computational Center” of Saint Petersburg State University (project no. 110-7198-609) for providing computing resources and data storage.

